# A diagnostic gap to fill: Development of molecular tools to distinguish the plant-parasitic nematode species *Heterodera carotae* and *Heterodera cruciferae*

**DOI:** 10.1101/2024.08.21.608931

**Authors:** Didier Fouville, Marine Biget, Josselin Montarry, Sylvain Fournet, Eric Grenier

**Affiliations:** IGEPP, INRAE, Institut Agro, Univ Rennes, Le Rheu, France

**Keywords:** molecular diagnostic methods, real-time PCR, cyst nematodes, microsatellite loci, sibling species

## Abstract

The carrot-parasitic nematode, *Heterodera carotae*, is an important pest causing qualitative and quantitative production losses across several countries worldwide and must be carefully identified and monitored. Indeed, in culture crops, *Heterodera carotae* and *Heterodera cruciferae*, two genetically closely related species from the Goettingiana group, can be found in mixture and are virtually morphologically non-recognizable. However, only *H. carotae* is able to develop on carrot. To monitor carrot production and set up suitable *H. carotae* control plans, simple and reliable diagnostic methods to detect and identify these two species are needed. In this study, we developed and successfully tested two real-time PCR protocols (SYBR Green singleplex and TaqMan multiplex real-time PCRs) against both sister species: *H. carotae* and *H. cruciferae*. Using two specific sets of primers targeting the two species, which were designed from sequences containing microsatellite loci, we managed, whatever the methods, to distinguish the two species. Moreover, we showed a higher specificity using the TaqMan assays (test on cysts) but a better sensitivity (tests on J2 juvenile larvae) with the SYBR Green assays. In this study, we thus proposed two useful and reliable diagnostic tools to enable farmers and scientists to rapidly detect two plant-parasitic nematodes and supervise agricultural management.

## 1. INTRODUCTION

The carrot cyst nematode, *Heterodera carotae* [1], is a plant-parasitic nematode showing an extremely narrow host-range, as it is able to attack and multiply only on wild and cultivated carrots (*Daucus carotae*), and incidentally on a wild herbaceous plant, *Torilis leptophylla* [2]. It is a polyvoltine species able to produce up to four successive generations during a growing season in favourable environmental conditions [3]. *Heterodera carotae* is a major issue causing qualitative and quantitative losses for carrot production [4, 5]. It is responsible for important carrot decline in several countries worldwide: it was reported in the USA, in Canada, in Mexico, in South-Africa, in India and in Europe [3, 5, 6, 7].

The ban of the last effective chemical nematicides (e.g. 1,3-dichloropropene), which were not specifically pointed at one species, needs to explore and find alternative control solutions targeting only the problematic parasitic species. This statement implies to be able to clearly identify the different nematodes at the species level for future carrot cyst nematode control plans. However, *H. carotae* is easily confused with its sister species *H. cruciferae* and are morphologically indistinguishable [8]. These two species belong to the Goettingiana genetic group [9] but differ by their host range, *H. carotae* being restricted to carrots and *H. cruciferae* to brassicas and a few species of Lamiaceae [10]. Because these two species could easily co-occur in mixed or rotated vegetable fields, and because host range tests are laborious and time-consuming, it is critical to develop suitable molecular tools able to distinguish these two species.

Madani et al. [11] developed molecular primers in the mitochondrial *cox1* gene which allowed specific identification of *H. carotae* populations parasitizing carrot from Canada and Italy. However, adding other *H. carotae* populations (from Mexico, Switzerland, France and Italy) and some *H. cruciferae* populations (from the USA and Russia), Escobar-Avila et al. [6] showed that using the different available molecular tools (including *cox1*), it was not possible to make an accurate differentiation of *H. carotae* from *H. cruciferae*. More recently and at a larger scale, Huston et al. [12] explored the utility of standard gene sequence barcodes for the delineation of species in the genus *Heterodera*. Using a database of 2,737 sequences of the four commonly used molecular markers in this genus (18S, ITS, 28S and *cox1*), their results showed that the mitochondrial cytochrome oxidase subunit one (*cox1*) region should be favoured for basic diagnostic purposes. Sequences of *cox1* allowed to distinguish all *Heterodera* species pairs, with just one case of a weak and one case of an inadequate pair delineation; the latter being the *H. carotae* / *H. cruciferae* pair [12].

Recently, Gautier et al. [7] developed a set of polymorphic microsatellite markers to explore the structuration of the genetic diversity of *H. carotae* populations at the European spatial scale and highlighted two distinct genetic clusters. Using the same set of markers at a smaller spatial scale in France, Esquibet et al. [13] demonstrated a high level of genetic divergence between *H. carotae* and *H. cruciferae* individuals and showed that some field populations were mixed populations containing both species. Microsatellites are highly variable neutral markers and most popular for geneticists. Gamel et al. [14] showed that microsatellite loci are also of interest to develop diagnostic markers and successfully developed a triplex real-time PCR assay for the detection of the cyst nematode species *Globodera pallida, Globodera rostochiensis* and *Heterodera schachtii*. It should be emphasized that in this case it is not the number of repeats of the microsatellite motifs that is considered for the diagnostic but instead the phylogenetic signal existing in the flanking regions of the microsatellites motif that allowed the design of species-specific primers and probes [14].

The aim of the present study was therefore to investigate the usefulness of about a hundred *H. carotae* microsatellite loci produced in Gautier et al. [7], and to develop and validate real-time PCR tools for the diagnosis of *H. carotae* and *H. cruciferae*. These novel molecular tools will allow scientists and farmers to time-monitor pest infection of the carrot crops and better supervise research and agricultural management.

## 2. MATERIALS & METHODS

### 2.1 Nematode populations

In total, 12 populations of *Heterodera carotae* (*Hca*), 4 populations of *Heterodera cruciferae* (*Hcr*), 1 population of *Heterodera goettingiana (Hgo)*, 1 population of *Heterodera avenae (Ha)* and 1 population of *Heterodera schachtii (Hs)* were included in this study (Table 1). All those 19 *Heterodera sp*. populations were obtained in our laboratory after multiplication on their specific hosts (carrot (*Hca*), rapeseed (*Hcr*), pea (*Hgo*) wheat (*Ha*) and beet (*Hs*)). Morphological, morphometric or host range criteria were used to identify and confirm the species status. Most of the *H. carotae* populations have been previously analyzed and clustered into two major phylogenetic clades [7].

**Table 1:** Nematode populations and species used in this study. Genetic clusters are indicated according to Gautier et al. (2019). NA: Not Applicable.

| Nematode species | Population name | Genetic cluster | Sampling site |
| --- | --- | --- | --- |
| *H. carotae* | Hca-0101 | Cluster 1 | Ain, France |
| *H. carotae* | Hca-1303 | Cluster 1 | Bouches du Rhône, France |
| *H. carotae* | Hca-4402 | Cluster 2 | Loire-Atlantique, France |
| *H. carotae* | Hca-ZAP | Cluster 1 | Italy |
| *H. carotae* | Hca-FU | Cluster 1 | Switzerland |
| *H. carotae* | Hca-02 | Cluster 2 | Aisne, France |
| *H. carotae* | Hca-2902 | Cluster 2 | Finistère, France |
| *H. carotae* | Hca-3001 | Cluster 2 | Gard, France |
| *H. carotae* | Hca-5006 | Cluster 2 | Manche, France |
| *H. carotae* | Hca-5601 | Cluster 2 | Morbihan, France |
| *H. carotae* | Hca-7201 | Cluster 2 | Sarthe, France |
| *H. carotae* | Hca-8401 | Cluster 2 | Vaucluse, France |
| *H. cruciferae* | Hcr-2201 | NA | Côtes d’Armor, France |
| *H. cruciferae* | Hcr-3501 | NA | Ille et Vilaine, France |
| *H. cruciferae* | Hcr-5001 | NA | Manche, France |
| *H. cruciferae* | Hcr-8601 | NA | Vienne, France |
| *H. goettingiana* | Hg-1701 | NA | Charente Maritime, France |
| *H. avenae* | Ha-Moulin | NA | Vendée, France |
| *H. schachtii* | Hs-AM02 | NA | Marne, France |

### 2.2 Microsatellite markers

Microsatellite markers were developed by Gautier et al. [7] according to the procedure described by Malausa et al. [15]. Briefly, purified DNA of 12 *H. carotae* populations (extracted from 100 cysts per population) were pooled and used to construct DNA libraries enriched in microsatellite motifs. From the NGS sequencing (454 pyrosequencing, ROCHE Diagnostics) of those libraries, 199 loci were identified as amplifiable targets (sequences with microsatellite motifs and successfully defined primers using Primer3 software). An additional sorting step discarding all tri-nucleotide motifs and the di-nucleotide motifs with less than 6 repetitions as well as small size loci (below 92 bp) led to a dataset of 95 loci. Among them, Gautier et al. [7] identified 13 polymorphic microsatellite markers usable in population genetic studies but in the present study we went back to the list of 95 loci and used them first in a conventional PCR screen in order to discard the ones showing bad or cross species amplifications.

### 2.3 DNA extraction of nematodes

Concentrated DNA solutions were produced for the two species of interest (*Hca* and *Hcr*) and others non-target *Heterodera* species (*Hgo, Ha, Hs*) using QIAamp DNA mini kit (Qiagen - protocol “DNeasy Blood and tissue”). Ten or 100 cysts were crushed into the Qiagen lysis buffer, DNA were eluted in 100µL of the Qiagen elution buffer according to the manufacturers’ instructions. DNA extracted from 100 cysts were used at 1/10 concentration to ensure a similar range of concentrations with the other 10-cyst extracted DNA samples. The quality of each DNA extraction was checked by successful amplification of the rDNA ITS fragment. To further test the sensibility of the real-time PCR tools, DNA was extracted from second-stage juveniles (J2) of the two target species *(Heterodera carotae* - Hca 1303 (genetic cluster 1), Hca 5006 (genetic cluster 2) and *Heterodera cruciferae* - Hcr 2201, Table 1). DNA was extracted from different numbers of J2 (one, two, five, ten and fifty). The detailed procedure for J2 DNA extractions is detailed in Gamel et al. [14]. Briefly, the freshly hatched J2 were transferred into a microtube containing a home-made lysis buffer. J2 were crushed using a Tissulyser II (Qiagen) and glass beads placed into the microtubes. Microtubes containing the crushed larvae were incubated at 55°C for 1h and then 95°C for 10 min to inactivate the proteinase K present in the lysis buffer. All the DNA extracts (cysts and larvae) were stored at -20 °C until molecular processing.

### 2.4 SYBR and TaqMan PCRs and validation

Primers and probes were designed from NGS sequence data using the Geneious software (version 8.0.5; Biomatters, http://www.geneious.com). The real-time PCR amplification parameters (SYBR and TaqMan methods) are presented in Tables 2 and 3. Real-time PCR reactions were performed on a LightCycler^®^ 480 instrument (Roche) using a specific master mix supplied by Roche Diagnostic. Specific master mixes LightCycler 480 Probes Master /Ref.: 04707494001 (Roche) and LightCycler 480 SYBR Green Master /Ref.: 04887352001 (Roche) were used to perform TaqMan and SYBR Green analyses, respectively. On the LightCycler^®^ 480 software, the automatic analysis option (Abs Quant/2nd derivative Mac) was selected to determine the cycle threshold (Ct). Real-time PCR assay has been tested in triplicate reactions to test the reproducibility of the measurement. Negative controls (water) were added to all the real-time PCR experiments. For the efficiency tests, four-fold serial dilutions of the targeted species (*H. carotae* and *H. cruciferae*) DNA were produced at known concentrations (2, 1, 0.5, 0.25 ng/µL).

**Table 2:**
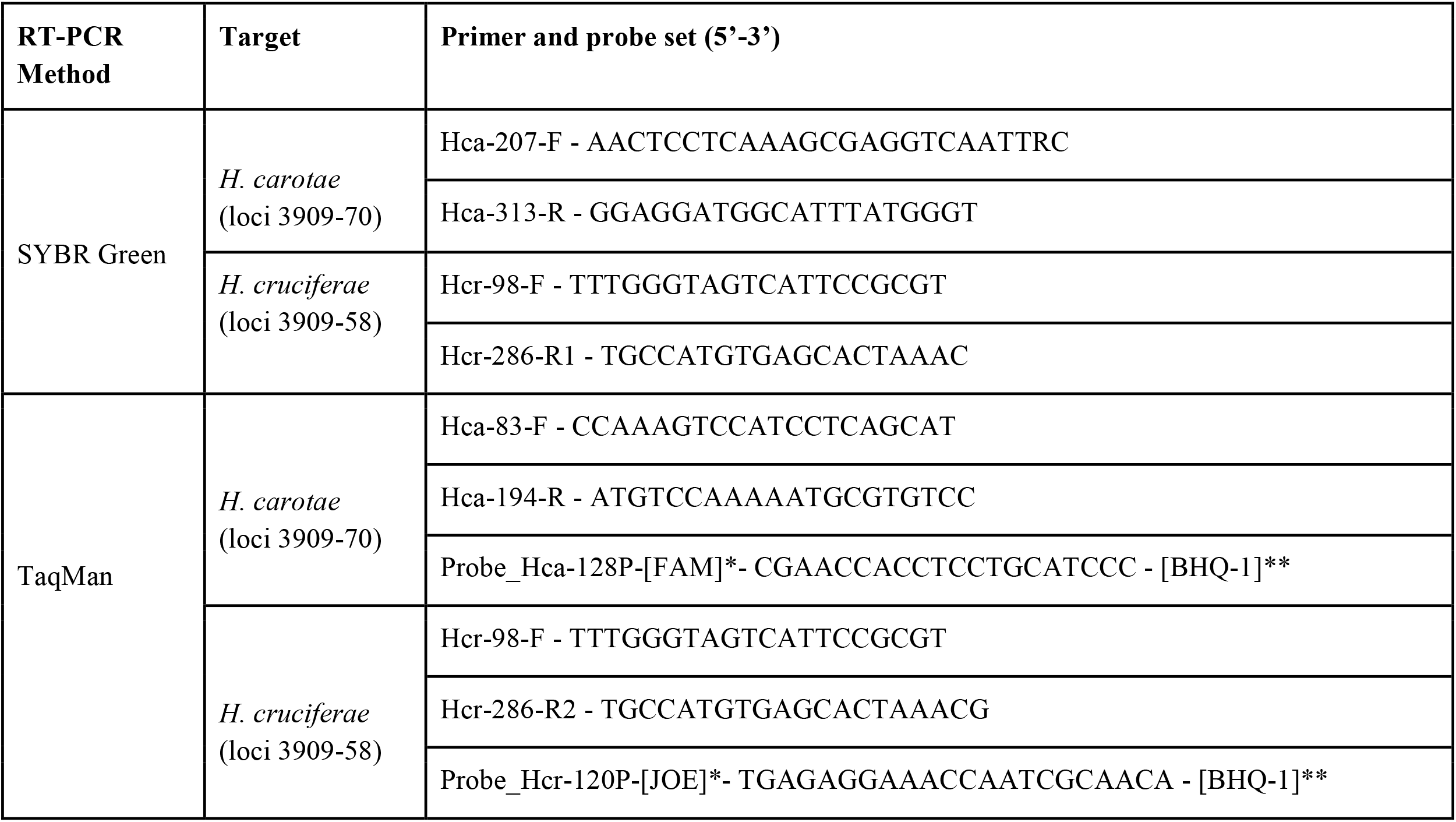
Primer and probe sets designed for molecular analyses. Note that amplicon size will slightly vary from one individual to the other due to variation in length of the microsatellite motif. The average amplicon size for both *H. carotae* pairs of primers is about 100 bp while amplicon size for both *H. cruciferae* pairs of primers is about 200 bp. * reporter; ** quencher.

**Table 3:**
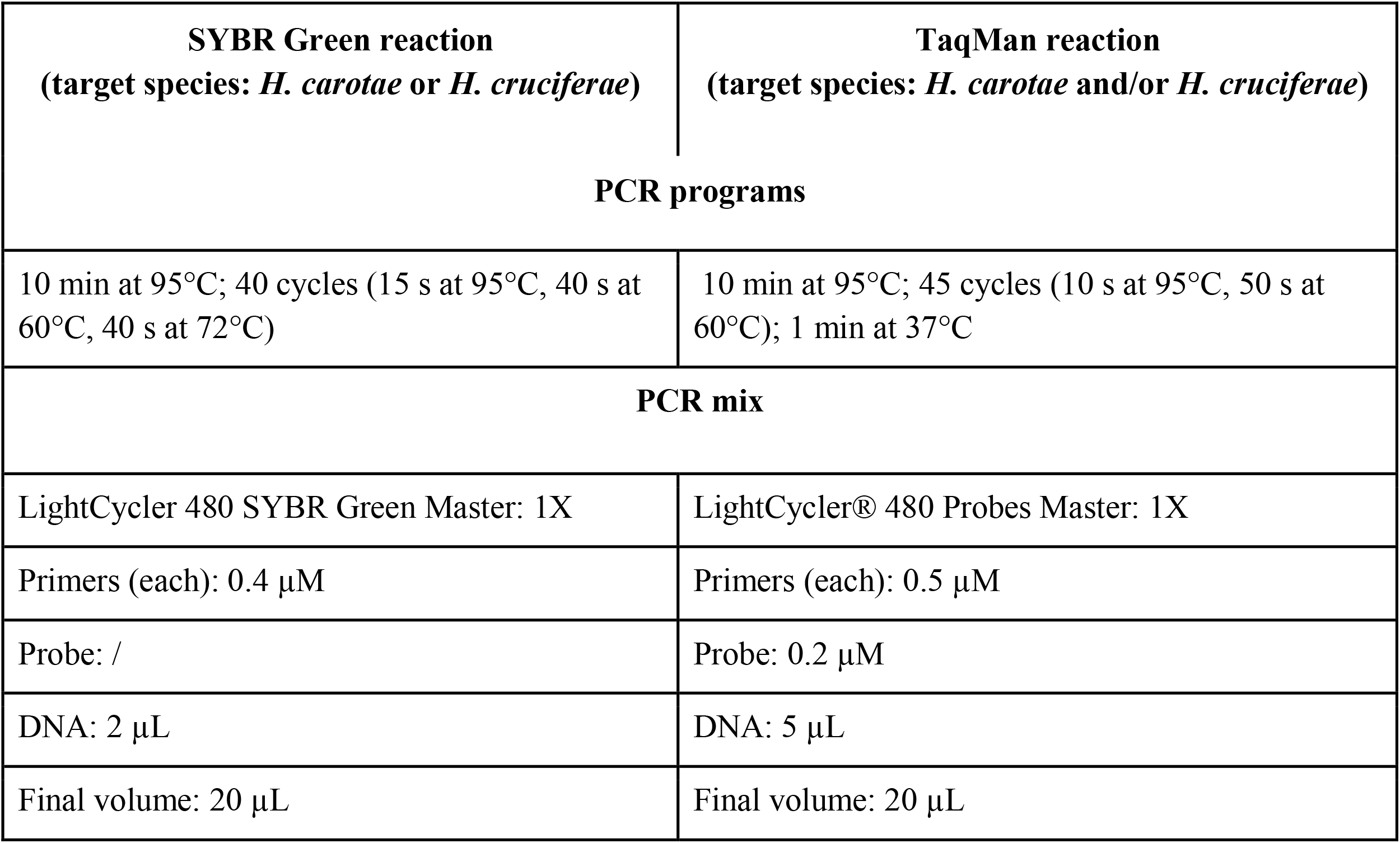
Mix composition and amplification programs for the two real-time PCR methods used in this study: SYBR Green and TaqMan RT-PCR.

| SYBR Green reaction<br>(target species: *H. carotae* or *H. cruciferae*) | TaqMan reaction<br>(target species: *H. carotae* and/or *H. cruciferae*) |
| --- | --- |
| PCR programs |  |
| 10 min at 95°C; 40 cycles (15 s at 95°C, 40 s at 60°C, 40 s at 72°C) | 10 min at 95°C; 45 cycles (10 s at 95°C, 50 s at 60°C); 1 min at 37°C |
| PCR mix |  |
| LightCycler 480 SYBR Green Master: 1X | LightCycler® 480 Probes Master: 1X |
| Primers (each): 0.4 µM | Primers (each): 0.5 µM |
| Probe: / | Probe: 0.2 µM |
| DNA: 2 µL | DNA: 5 µL |
| Final volume: 20 µL | Final volume: 20 µL |

## 3. RESULTS

In this study, we developed and validated rapid and efficient real-time PCR assays to distinguish sister species often co-occuring in mixed or rotated vegetable fields producing carrots: *Heterodera carotae* (*Hca*) and *Heterodera cruciferae* (*Hcr*). To do so, we first screened 95 microsatellite loci using conventional PCR on 11 populations (including five *H. carotae* and three *H. cruciferae*) to identify the best candidates. Then, we developed and tested the selected loci using two different methods, the simplex real-time PCR (the SYBR Green tool) and the duplex real-time PCR (the TaqMan tool); at two levels of sensitivity, the cyst and the larvae levels.

### 3.1 Primer specificity revealed during first screen of the microsatellite loci

During the first screen by conventional PCR we identified 42 microsatellite loci showing either multiple PCR products in *Hca* populations or no amplification with the tested *Hca* and *Hcr* DNAs (data not shown). These loci were discared and analysis was conducted only on the 53 remaining (Table S1). The diagnostic specificity, sensitivity and the relative accuracy criteria were assessed by calculating the number of positive and negative agreements and deviations compared to the status of the sample as described in the EPPO standard protocol for test validation PM7/98 (2) [16]. The experiment was conducted using only a subset of *Hca* (5/12) and *Hcr* (3/4) populations. Five microsatellite loci showing diagnostic specificity (i.e. the ability to not detect two different species) of 100% and 48 loci showing diagnostic sensitivity (i.e. the ability to detect all populations from a target species) of 100% were identified (Table S1). Only two microsatellite loci showed a relative accuracy of 100%: loci 3909-70 for detection of *H. carotae* and loci 3909-58 for detection of *H. cruciferae* (Table S1). These two microsatellite loci were therefore selected for further analysis using real time PCR and using a larger set of *H. carotae* and *H. cruciferae* populations.

### 3.2 Simplex real-time PCR assays on nematode cysts

The two sets of specific primers 3909-70 and 3909-58 were individually tested against the targeted (*Hca* and *Hcr*) and non-targeted nematode species (*Hgo, Ha* and *Hs*) using the SYBR Green tool. The Ct values of each triplicate per population, whatever the marker, were close enough to be confident in the results and the primer performances (Table 4). The Ct values obtained with the 3909-70 marker specific to *Hca* ranged from 21.5 to 27.2 for the 12 *Hca* populations, irrespective of the cluster to which they belonged, and from 26.1 to 30.0 for the 4 *Hcr* populations including the Hcr-3501 population, for which Ct values were closed to those obtained for the *Hca* populations (Table 4). However, the Tm values obtained for the *Hcr* populations allowed to clearly discriminate between the two targeted populations with at least 1.2°C difference (Table 4). The Ct values obtained for the non-targeted species and the no template controls (NTC) were higher than 35, which is usually taken as the limit of detection (LOD). The Ct values obtained with the 3909-58 marker specific to *Hcr* ranged from 21.3 to 25.9, values much lower than those obtained for *Hca* and the non-targeted species (Table 4). Indeed, most of the *Hca* Ct values, including those of the non-targeted species and NTC samples, were higher than 31.9 and mainly higher than the LOD. It seems that *Hca* populations from cluster 1 were less well recognized by this marker compared to *Hca* populations from cluster 2 as 3 out of 4 cluster 1 populations are not detected or showed Ct values > 35 while only 3 out of 8 cluster 2 populations showed Ct values > 35.

**Table 4:** Specificity test of the *Hcr* (3909-58) and *Hca* (3909-70) markers using SYBR Green real time PCR reaction on DNA from *H. carotae* (12 populations), *H. cruciferae* (4 populations), *H. goettingiana, H. avenae* and *H. schachtii*. Cycle threshold (Ct) and melting temperature (Tm) are indicated for both markers, each population and each replicate. ND: Not Detected.

|  | Ct observed for<br>marker 3909-58 |  |  | Tm observed for<br>marker 3909-58 |  |  | Ct observed for<br>marker 3909- 70 |  |  | Tm observed for<br>marker 3909- 70 |  |  |
| --- | --- | --- | --- | --- | --- | --- | --- | --- | --- | --- | --- | --- |
| **Hca-FU** | ND or > 35 |  |  | ND |  |  | 23,8 | 23,6 | 23,7 | 79,0 | 79,0 | 79,1 |
| **Hca-ZAP** | 32,4 | 33,9 | 33,0 | 78,5 | 78,4 | 78,4 | 26,7 | 26,4 | 26,5 | 79,0 | 79,1 | 79,1 |
| **Hca-0101** | ND or > 35 |  |  | ND |  |  | 24,6 | 24,6 | 24,5 | 79,1 | 79,1 | 79,1 |
| **Hca-02** | 32,9 | 32,4 | 32,5 | 78,6 | 78,6 | 78,7 | 24,3 | 24,4 | 24,3 | 78,8 | 78,9 | 78,9 |
| **Hca-1303** | ND or > 35 |  |  | ND |  |  | 22,7 | 22,8 | 23,6 | 79,0 | 79,0 | 79,2 |
| **Hca-2902** | ND or > 35 |  |  | ND |  |  | 27,2 | 27,2 | 27,2 | 78,9 | 78,9 | 79,0 |
| **Hca-3001** | ND or > 35 |  |  | ND |  |  | 23,3 | 23,4 | 23,6 | 78,9 | 78,9 | 78,8 |
| **Hca-4402** | ND or > 35 |  |  | ND |  |  | 23,3 | 23,2 | 23,1 | 78,9 | 79,0 | 79,0 |
| **Hca-5006** | 34,7 | 33,9 | 35,2 | 78,9 | 78,8 | 78,9 | 22,9 | 22,9 | 22,8 | 79,1 | 79,1 | 79,2 |
| **Hca-5601** | 31,9 | 32,5 | 32,6 | 78,7 | 78,7 | 78,2 | 21,6 | 21,6 | 21,5 | 79,2 | 79,2 | 79,2 |
| **Hca-7201** | 34,9 | 36,5 | 34,8 | 75,6 | 75,2 | 75,6 | 22,8 | 22,7 | 22,8 | 79,2 | 79,1 | 79,2 |
| **Hca-8401** | 33,9 | 34,4 | 41,3 | 83,0 | 83,0 | 83,0 | 23,7 | 22,5 | 23,5 | 79,1 | 79,1 | 79,1 |
| **Hcr-2201** | 23,7 | 23,6 | 23,6 | 78,8 | 78,8 | 78,8 | 28,5 | 28,2 | 28,3 | 77,5 | 77,6 | 77,6 |
| **Hcr-3501** | 22,0 | 22,3 | 22,3 | 78,7 | 78,7 | 78,7 | 26,2 | 26,2 | 26,1 | 77,5 | 77,5 | 77,5 |
| **Hcr-5001** | 25,7 | 25,8 | 25,9 | 78,7 | 78,8 | 78,7 | 30,0 | 29,9 | 29,8 | 77,5 | 77,5 | 77,6 |
| **Hcr-8601** | 21,3 | 21,4 | 21,3 | 78,6 | 78,7 | 78,7 | 27,9 | 27,8 | 27,9 | 77,6 | 77,6 | 77,6 |
| **Hg-1701** | 35,7 | 34,2 | 34,7 | 78,8 | 78,9 | 78,9 | > 35 |  |  | ND |  |  |
| **Ha-Moulin** | > 35 |  |  | ND |  |  | > 35 |  |  | ND |  |  |
| **Hs-AM02** | > 35 |  |  | ND |  |  | > 35 |  |  | ND |  |  |

### 3.3 Duplex Real-Time PCR Assays on nematode cysts

Two sets of specific primers and probes (Table 2) were tested against the targeted (*Hca* and *Hcr*) and non-targeted nematode species (*Hgo, Ha* and *Hs*) using the TaqMan tool (Figure 1 and Table S2). This method, using specific primers and probes, allowed the multiplexing of the two analyses (*Hca* and *Hcr* detection assays) in a single reaction. Standard curves showed a very high level of sensitivity of the method for the two targeted species (efficiency values reaching 2.081 and 2.182 for the *Hca* and the *Hcr* detection respectively; Figures 1B and 1C). No signal was detected in the non-targeted species and NTC samples. Multiplex analysis results clearly clustered the two targeted populations depending on their specific fluorescence wavelength (blue dots for *Hca* and green dots for *Hcr*) and dissociated the non-targeted species as well as the controls (grey dots) (Figure 1A). Moreover, Ct values obtained for the two targeted species were similar to the ones observed for the targeted species in previous simplex real-time PCR assays (Table S2 and Table 4). *Hca* populations showed lower Ct values, irrespective of the cluster to which they belonged, compared to *Hcr* populations (Table S2). Therefore, the use of the multiplex analysis associated with the TaqMan tool allowed precisely distinguishing the two sister species in a single run.

**Figure 1:**
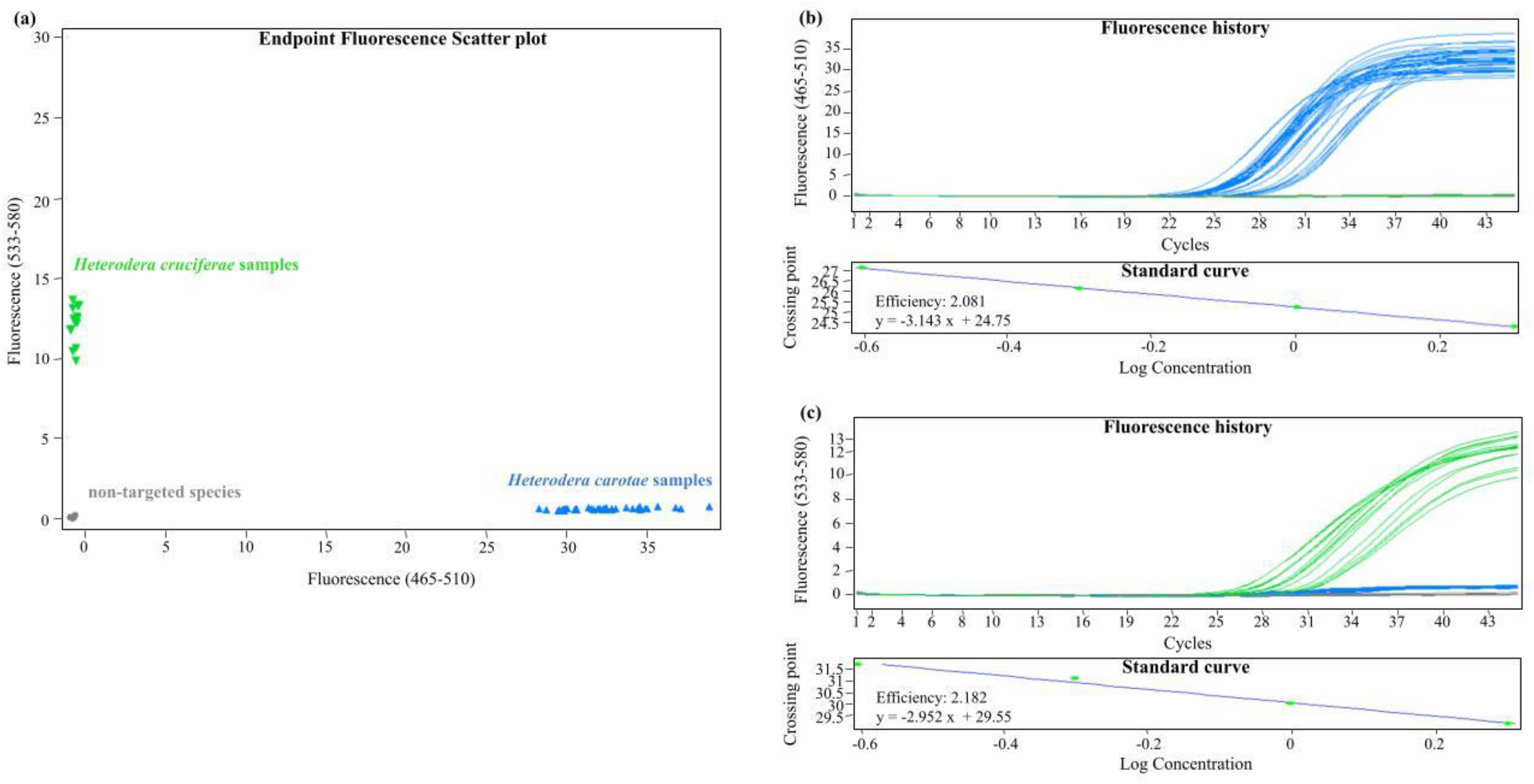
Graphical outputs of genotyping results in the TaqMan assay. In A, the allelic discrimination plots of end-point fluorescent TaqMan PCR data of the 2 probes analyzed in 19 populations of *Heterodera* including 12 *H. carotae*, 4 *H. cruciferae*, 1 *H. goettingiana*, 1 *H. avenae* and 1 *H. schachtii*. FAM fluorophore (x -axis) is associated with the probe for *Hca* diagnostic amplification (blue dots), while JOE (y -axis) labels the *Hcr* diagnostic amplification (green dots). Non-target species are indicated by grey dots. In B, the amplification curves (top) and a standard curve (bottom) for detection of *Hca* diagnostic marker. In C, the amplification curves (top) and a standard curve (bottom) for detection of the *Hcr* diagnostic marker.

### 3.4 Detection level sensitivity of the two methods on larvae samples

We assessed the analytical sensitivity of our two diagnostic tools (simplex and duplex real-time PCRs) using DNA extracted from different numbers of J2 (one, two, five, ten and fifty) of two populations of *Hca* and one population of *Hcr*. All these DNA extractions were each performed in a total volume of 100 µL but either 2 µL or 5 µL were used depending on the diagnostic tool tested. The LOD was considered as samples giving positive results (Ct values < 35) in all replicates. No signal was detected in the NTC samples (data not shown). Using 2 µL of DNA extracted from 50 J2 in our simplex *Hca* or *Hcr* real-time PCRs resulted in Ct values < 35 in all replicates (i.e. 27.9 and 28.9 for the Hca-1303 and Hca-5006 populations respectively; 32.7 for the Hcr-2201 population) (Figure 2). As PCRs were conducted using a 1/50 dilution (2 µL out of 100 µL) we can assume that 1 J2 (50 J2 diluted 1/50) is the LOD for our simplex reactions. We were unable to detect precisely the LOD for our duplex TaqMan reaction as using 5 µL of DNA extracted from 50 J2 in our TaqMan real-time PCR did not result in Ct values < 35 for all tested populations (Figure 3). However, by linear extrapolation and considering the dilution factor linked to our TaqMan reaction we can predict that the LOD for the TaqMan duplex reaction should be around 100 J2, meaning that still a single cyst should be enough for an accurate diagnostic.

**Figure 2:**
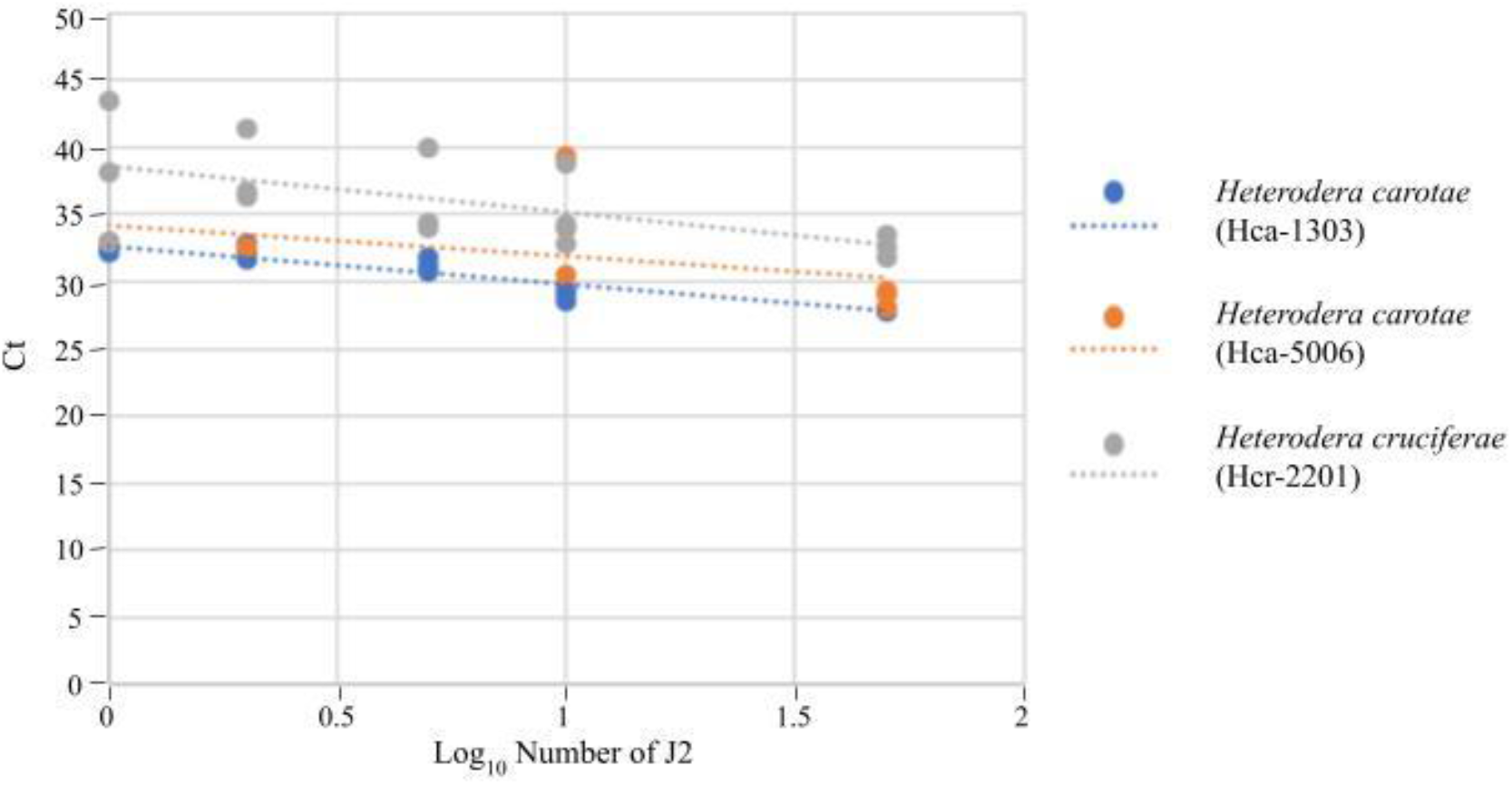
SYBR Green diagnostic sensitivity. Cycle threshold values obtained by testing a range of juveniles (J2) of *H. carotae* Hca-1303 and Hca-5006 and *H. cruciferae* Hcr-8601 target species in simplex SYBR Green real time PCR reaction. The number of J2 is expressed on a logarythmic scale.

**Figure 3:**
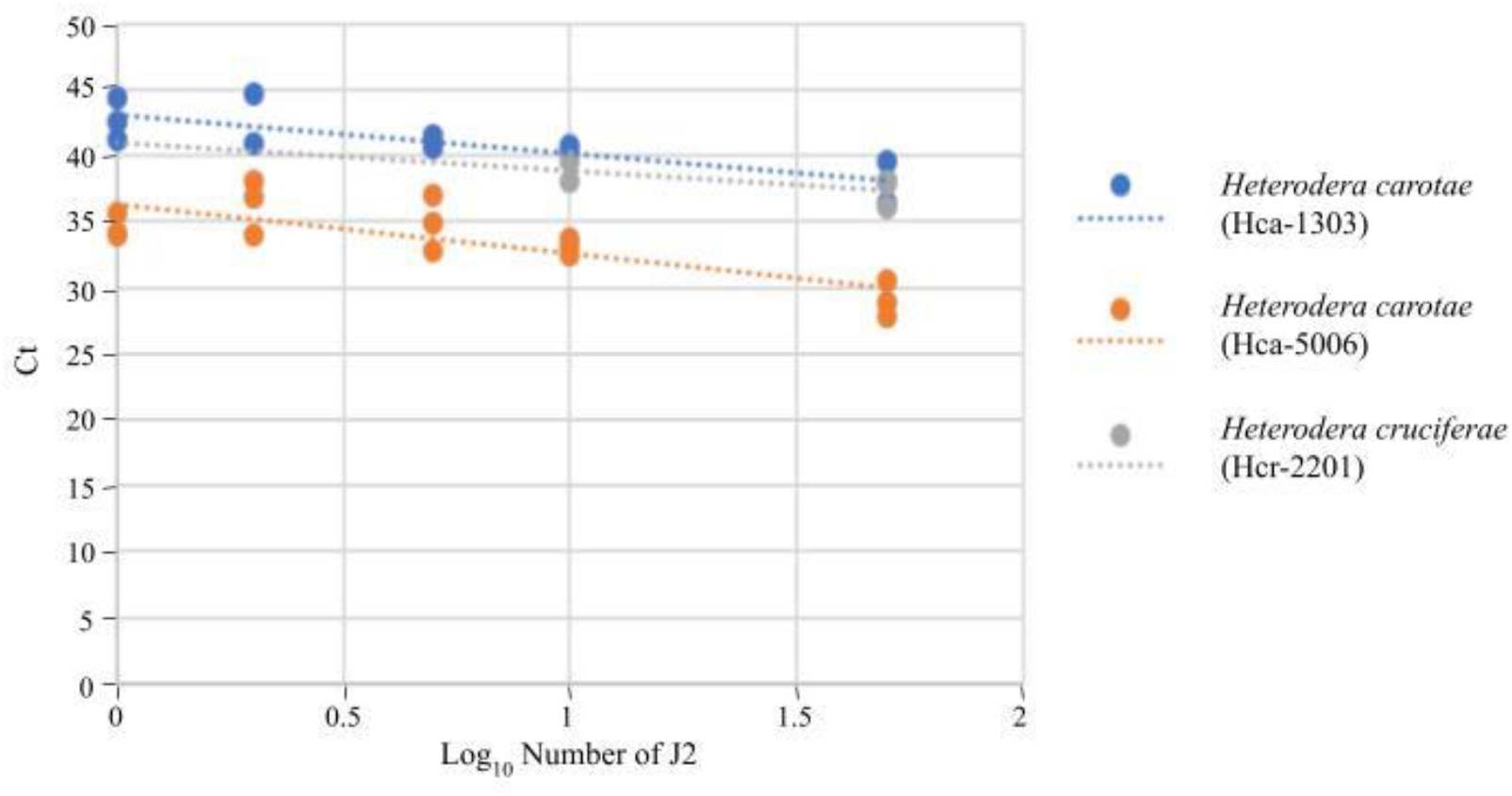
TaqMan diagnostic sensitivity. Cycle threshold values obtained by testing a range of juveniles (J2) of *H. carotae* Hca-1303 and Hca-5006 and *H. cruciferae* Hcr-8601 target species in duplex TaqMan real time PCR reaction. The number of J2 is expressed on a logarythmic scale.

## 4. DISCUSSION

### 4.1 A diagnotic tool for a better management of the *H. carotae* issue

*Heterodera carotae* is a major phytosanitary problem for the carrot sector, as yield losses could reach 80% [4, 5]. Since the withdrawal of the last effective chemical nematicide and because there are no carrot cultivars resistant to *H. carotae*, the European carrot producers have modified their rotation schemes. Among plants introduced into those rotation schemes, many cruciferous vegetables, such as cauliflower, cabbage or broccoli, are non-host plants for *H. carotae* but host plants for *H. cruciferae*. Moreover, some plants from the Brassica family which were used as biofumigant plants, such as the mustards, could also be hosts of *H. cruciferae*.

The situation became thus more and more favorable to the co-occurrence of both species in the same field as shown in France [13], and makes the nematological analyses – which so far only indicate the number of cysts and larvae belonging to the Goettingiana group - inappropriate for making adequate control decisions. The molecular tools developed in the present study allow now to distinguish these two nematode species. Following the cysts extraction from the soil, the analytical laboratories would be able to used a molecular tool (i.e. two conventional PCR, two SYBR Green PCR or a unique Taqman PCR) on a DNA sample obtained from pools of cysts or on several DNA samples obtained from individual cysts in order to estimate the proportion of both species in the sampled field.

### 4.2 An improvement over current diagnostic tools and novelty

Many molecular tools have been developed and tested over the past decades to identify nematode species of interest in routine: conventional PCR (SCAR), random amplification of polymorphic DNA (RAPD), amplified fragment length polymorphism (AFLP) and PCR amplification and sequencing of a specific gene [17]. All these methods showed advantages but important drawbacks in terms of sensitivity (e.g. samples with low amount of DNAs), most of them being time-consuming and laborious to set up in a classic laboratory. Real-time PCR method allows rapid, specific and sensitive detection of a variety of nematodes including cyst nematodes [17]. In recent studies, different research teams have reported successfully identification of *Heterodera glycines* from other related and closed species *H. trifolii* and *H. schachtii* using real-time PCR and new primers targeted the ITS region or CLE genes segment [18, 19]. Recently, Subbotin et al. [20] also developed and validated a multiplex PCR tool (TaqMan) to detect *Heterodera mediterranea* using an universal *hsp90* gene primer set and a specific primer set against *H. mediterranea*. The authors showed a high level of sensitivity using the specific primer set: detection of a 0.03-dilution of J2 or one J2 for *in vitro* and field sample assays respectively [20].

Although several studies have been conducted using real-time PCR to identify and quantify various cyst nematodes with very sensitive and precise methods, to our knowledge, only Gamel et al. [14] have developed molecular markers using microsatellite loci that enable precise identification of different and closely related species. They showed high sensitivity (LOD of 5 J2) and specificity in detecting *H. schachtii* among 11 non-target species including the *Heterodera betae* sibling species. In the same study they also investigated the usefulness of such microsatellite loci for the development of diagnostic tools to distinguish *Globodera pallida* and *G. rostochiensis*. They reported an efficient separation of these two species from their respective sister species (i.e. *G. mexicana* for *G. pallida* and *G. tabacum* for *G. rostochiensis)* with a sensitivity threshold of a single J2 [14]. In recognition of its value, the EPPO standard diagnostic protocol for *G. rostochiensis* and *G. pallida* PM 7/40 (5) [21], now also recommends the Gamel et al. [14] PCR based tool. In this study, we confirmed once again the interest of microsatellite loci to identify species-specific polymorphisms and design novel diagnostic tool. The procedure we followed was similar to the one described in Gamel et al. [14], except that for *H. carotae* we started from less amplifiable loci (199), while 481 and 354 were used as starting material for *G. rostochiensis* and *H. schachtii*, respectively. Nonetheless, the approach was successful as we specifically detected our two targeted species, recorded a one-J2 threshold for our simplex reactions and managed to separate the two sister species in a duplex reaction.

### 4.3 Target and non-target populations used and limits remaining to be investigated

We used a total of 12 *Hca* populations representing the genetic diversity present in our collection and expected to cover a large part of the *Hca* European genetic diversity. Detection using either the TaqMan or SYBR Green 3909-70 tools were always achieved in a way irrespective of the genetic cluster to which the population belonged. Other molecular investigations will be required to clearly assign *Hca* populations to one of the two main genetic clusters, but one can notice that using the SYBR Green 3909-58 tool (for detection of *Hcr*), we were able to better detect *Hca* populations from cluster 2 as 62.5% of these populations showed Ct values below the LOD while only 25% of the *Hca* populations from cluster 1 showed Ct values below the LOD.

Some limits remain to be investigated in future studies. The main one being the evaluation of these tools against further non-target species. *Heterodera carotae* and *H. cruciferae* belong to the Goettingiana group, which also contains the following species *H. goettingiana, H. urticae, H. persica* and the recently new species described in China *H. microulae* [22]. Even if the main species of this group were used in this study to assess the specificity, half of the species diversity known to date in the Goettingiana group remains to be investigated. Hence, it will be of interest to confirm the absence of cross-reaction of these new diagnostic tools with the remaining species *H. urticae, H. persica* and *H. microulae* and also with representatives of other plant parasitic species that can be found in carrot crop fields (i.e. *Pratylenchus, Radopholus* and *Meloidogyne*).

Also, it should be noticed that we have no idea if the four *Hcr* populations we used are representative of the genetic diversity in this species. We are confident that these four *Hcr* populations represent the diversity in France but further populations from other European countries will need to be tested to confirm the species specificity of the TaqMan assay. In terms of sensitivity, the LOD for the *H. carotae* SYBR Green assay was one single juvenile with a Ct value in between 27.9 and 28.9 for *H. carotae* and a Ct value of 32.7 for *H. cruciferae*. However, for the TaqMan assay, the LOD was not precisely determined and future investigations should extend the number of J2s tested up to 200 J2s and even one cyst in order to confirm the estimation of a LOD assumed in this study.

Finally, we have not evaluated the tools described here using DNA directly extracted from soil samples containing nematodes and further developments addressing this limit will clearly be also of interest. A TaqMan real time PCR was recently successfully applied to detect *Heterodera filipjevi* in field soils samples and showed a correlation (R^2^=0.82) between the numbers of this nematode found by conventional methods and the numbers quantified using the qPCR assay [23]. A DNA-based soil testing service operates in Australia using TaqMan assays and allowing a range of fungal and nematode plant pathogens to be quantified. In their experience of several years, the TaqMan assays were shown to overestimate the population density of plant nematodes compared to the traditional method, but showed a good relationship (R^2^=0.84) between nematode estimates obtained using the traditional and real time PCR methods [24].

### 4.4 Microsatellite loci as regions of interest to develop species-specific primers

In this study, we used microsatellite loci generated in the original objective to study the population genetics of the carrot cyst nematode *H. carotae*. Microsatellites have a well documented and recognized usefulness for population genetics studies, but a much less recognized usefullness for the development of diagnostic tools. However, given their large number in nematode genomes (up to 4480 in root-knot nematode genomes, Castagnone-Sereno et al. [25] and up to 22,600 in potato cyst nematode genomes, Eves-van den Akker et al. [26]) and given the existence of some phylogenetic signal in their flanking genomic regions, these markers appear to be an alternative to the regions classically used, such as ribosomal and mitochondrial DNA. First attempts to use microsatellites for molecular diagnostic of nematodes were conducted with *Heterodera schachtii, Globodera pallida* and *G. rostochiensis* [14]. Here, we have used species of cyst nematode (*H. carotae* and *H. cruciferae*) belonging to the Goettingiana group which is genetically distant from the Schachtii group in the *Heterodera* genus [9]. Furthermore, in the case of *H. carotae*, the sister species *H. cruciferae*, which is morphologically and molecularly indistinguishable [8], added a significant challenge that may have hampered the success of a molecular diagnostic tool. Unsurprisingly, we found very few loci after our initial selection that met our criteria for relative accuracy and we also observed cross-reactions using both SYBR Green markers even if a threshold Ct value in the case of marker 3909-58 or a threshold Tm value in the case of marker 3909-70 can be used to distinguish *Hca* from *Hcr*. Actually, we were fortunate to identify one locus specific to *H. carotae* and one specific to *H. cruciferae*, as neither of these two loci alone would have allowed easy and unambiguous identification.

### 4.5 Conclusion

Using sequences generated to identify microsatellite loci, we have successfully developed a SYBR Green and a Taqman method that can be used to easily distinguish the two geneticaly and morphologically closely related species *H. carotae* and *H. cruciferae*. We highlighted that these two real-time PCR tools were complementary, as the former showed a better sensitivity whereas the latter showed a better specificity. The increasing availability of genome data for a wide range of nematodes, together with methodological developments of *in silico* mining of microsatellites, allow for the easier characterization and use of these markers in the future. However, developing diagnostic tools for the diversity of nematode species encountered in crop fields has the makings of a never-ending effort, especially when considering extensive field surveys and description of the existing plant-parasitic nematode communities that is still a goal in terms of pest management. Thanks to the development of third generation sequencing platforms, the future of nematode molecular diagnosis may then rather lie in the use of long target sequencing than the development of species-specific tools. Genomic sequencing approaches have the potential to capture evolutionary and phylogenetic signal of species better than single locus analyses and should be able to deal better with mixed samples from complex natural environments. Some recent studies using long read sequencing of rDNA in nematodes have confirmed the interest and feasibility of such approach despite either some technical challenge related for example to the addition of MinION tails [27] or some ambiguous taxonomic assignation in multi-species samples [28].

## STATEMENTS AND DECLARATIONS

### Author contributions

DF and EG conceived the project. DF and EG, helped with MB, SF and JM, set up the methodology of the study. DF conducted lab experiments and DF and EG analyzed the data. DF and MB wrote the first draft of the paper. All the authors gave their final approval for publication.

## Acknowledgments

We thank the colleagues from the Anses and INRAE NemAlliance group (https://www.anses.fr/fr/content/nemalliance) for their expertise and help in the confirmation of the taxonomic status of the populations used in this study.

## FIGURES LEGEND

Table S1: Results obtained by conventional PCR for the selection of relevant loci for H. carotae and H. cruciferae detection by testing a range of target and non-target populations and calculating the diagnostic specificity, the diagnostic sensitivity and the relative accuracy.

Table S2: Specificity test of the Hca (3909-70) and Hcr (3909-58) markers using TaqMan real time PCR reaction on DNA from H. carotae (12 populations), H. cruciferae (4 populations), H. goettingiana, H. avenae and H. schachtii. Cycle threshold (Ct) are indicated for both markers, each population and each replicate. ND: Not Detected.

